# A comparative baseline of coral disease across the central Red Sea

**DOI:** 10.1101/2021.01.28.428586

**Authors:** Greta Smith Aeby, Amanda Shore, Thor Jensen, Maren Ziegler, Thierry Work, Christian R. Voolstra

## Abstract

The Red Sea is a unique environment for corals with a strong environmental gradient characterized by temperature extremes and high salinities, but minimal terrestrial runoff or riverine input and their associated pollution. Disease surveys were conducted along 22 reefs in the central Red Sea along the Saudi Arabian coast in October 2015, which coincided with a bleaching event. Our objectives were to 1) document types, prevalence, and distribution of coral diseases in a region with minimal terrestrial input, 2) compare regional differences in diseases and bleaching along a latitudinal gradient of environmental conditions, and 3) use histopathology to characterize disease lesions at the cellular level. Coral reefs of the central Red Sea had a widespread but a surprisingly low prevalence of disease (<0.5%), based on the examination of >75,750 colonies. Twenty diseases were recorded affecting 16 coral taxa and included black band disease, white syndromes, endolithic hypermycosis, skeletal eroding band, growth anomalies and focal bleached patches. The three most common diseases were *Acropora* white syndrome (59.1% of the survey sites), *Porites* growth anomalies (40.9%), and *Porites* white syndrome (31.8%). Over half of the coral genera within transects had lesions and corals from the genera *Acropora, Millepora* and *Lobophyllia* were the most commonly affected. Cell-associated microbial aggregates were found in four coral genera resembling patterns found in the Indo-Pacific. Differences in disease prevalence, coral cover, amount of heat stress as measured by degree heating weeks (DHW) and extent of bleaching was evident among sites. Disease prevalence was not explained by coral cover or DHW, and a negative relationship between coral bleaching and disease prevalence was found. The northern-most sites off the coast of Yanbu had the highest average DHW values but absence of bleaching and the highest average disease prevalence was recorded. Our study provides a foundation and baseline data for coral disease prevalence in the Red Sea, which is projected to increase as a consequence of increased frequency and severity of ocean warming.

## Introduction

Coral disease is a significant factor impacting coral reefs with localized disease outbreaks occurring worldwide [1,2]. The single most damaging disease outbreak, stony coral tissue loss disease, has been devastating coral reefs throughout the Florida Reef Tract since 2014 [3–5] and has spread to neighboring Caribbean regions [6]. Outbreaks of coral disease have increased through time [7] and have been linked to anthropogenic impacts such as overfishing [8], plastic pollution [9], dredging activities [10], terrestrial runoff [11,12], and increased ocean temperatures [13,14].

Coral reefs face different threats depending on their geographic region. The Red Sea is a unique environment for corals being a partially enclosed body of water with limited exchange with the Indian Ocean, low influxes of freshwater (~30mm/year) and high evaporation rates [15,16]. The majority of coral reefs around the globe live with temperatures usually not exceeding 29°C and salinities around 36 PSU [17]. In the Red Sea, temperature extremes are the norm with temperatures surpassing 32°C in the summer, around 18°C in the winter and with salinities 40 PSU or higher [18,19]. Yet, the Red Sea has extensive and healthy coral reefs with approximately 346 species of reef corals [20,21]. The Red Sea is characterized by natural north to south gradients of temperature, salinity and nutrient availability [22,23]. As example, in the far north, sea surface temperatures (SSTs) average 26 °C (± 1 °C) compared to 31.3 °C (± 1.1 °C) in the south [22]. Numerous bleaching events have occurred on coral reefs in the Red Sea which also show a latitudinal gradient in coral response. During the recent bleaching in 2015, Osman et al. [24] reported that degree heating weeks (DHW) surpassing the bleaching threshold of 4 (https://coralreefwatch.noaa.gov/product/5km/index_5km_dhw.php) occurred throughout the Red Sea, yet bleaching was restricted to the central and southern Red Sea becoming more severe to the south.

Although coral reefs in the Red Sea face temperature and salinity extremes typically not experienced by corals in other ocean basins, they also receive minimal terrestrial runoff or riverine input and their associated sedimentation, turbidity, and nutrient enrichment. Terrestrial runoff degrades local coral reefs [25] and contributes to increased severity and prevalence of coral diseases [10,26–28]. This creates a unique opportunity to examine coral health in an ecosystem with naturally high temperatures and salinities but minimal terrestrial pollution. We conducted coral disease surveys along the Saudi Arabian coast of the Red Sea in October 2015 which coincided with a bleaching event. Our objectives were to 1) document types, prevalence, and distribution of coral diseases in a region with minimal terrestrial input, 2) compare regional differences in diseases along a latitudinal gradient of environmental conditions and a gradient of bleaching response, 3) use histopathology to characterize disease lesions at the cellular level.

## Materials and methods

### Disease and bleaching surveys

Coral community structure, disease prevalence and bleaching extent was recorded at 22 sites along the Red Sea coast of Saudi Arabia between October 20 and November 9, 2015 (S1Table; Fig 1). At each site, divers surveyed two replicate 25m belt transects deployed end to end separated by approximately five meters. Corals were identified to the genus level along 25 x 1m belts with the exception of some taxa that are difficult to distinguish in the field. As such, *Favites* and *Dipsastraea* were combined, *Goniopora* and *Alveopora* were combined and *Lobophyllia* and *Symphyllia* were combined. Substrate characteristics (hard coral, soft coral, crustose coralline algae, macroalgae, rubble, sand) and bleaching (color loss) were documented by point-intercept method with the substratum underlying the tape measure recorded at 25 cm intervals and coral cover scored as bleached (pale to total color loss) or healthy. Coral lesions were assessed along wider 25 x 6 m belts. Gross lesions were classified into three lesion types including tissue loss, discoloration and growth anomaly, and nomenclature for lesions was based on the host genus affected and the lesion type (e.g. *Acropora* growth anomaly; [29]). Tissue-loss lesions were further classified based on the lesion size, shape, presence of predators, knowledge of what common predation marks look like and evidence of lesion progression based on degree of algal colonization onto the bare coral skeleton. Transect lengths, widths and numbers were modified as needed when constrained by dive limits.

**Fig 1.**
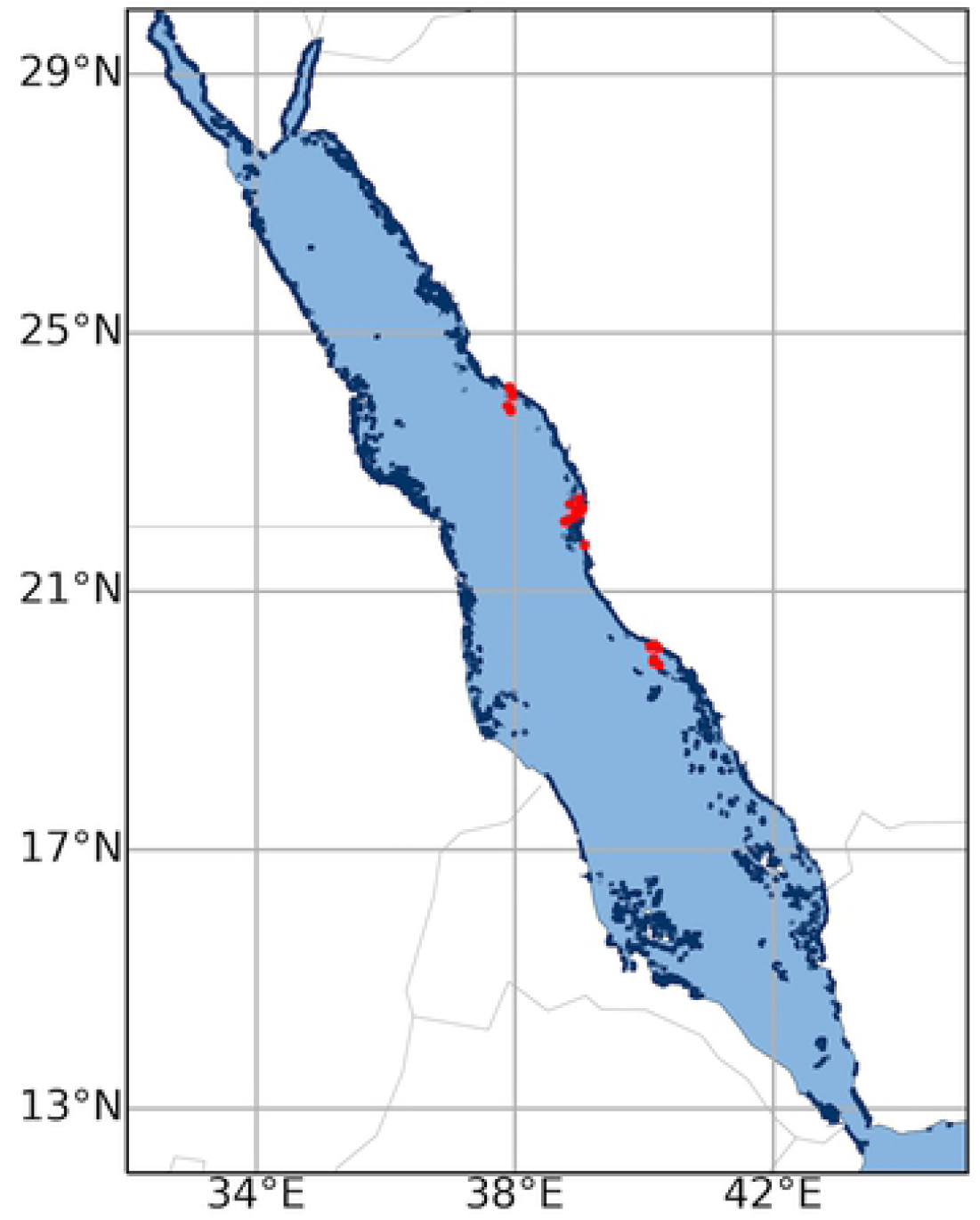
Sites surveyed for coral disease in three regions along the Saudi Arabian coast within the central Red Sea in Oct-Nov, 2015. Red dots indicate survey sites and blue dots indicate presence of coral reefs.

### Histopathology of coral lesions

Paired normal and lesion tissues of coral lesions encountered during surveys were sampled for histopathologic analysis to characterize host response and presence of organisms visible on light microscopy [30]. Fragments were fixed in 20% zinc formaldehyde-seawater immediately after the dive and processed for routine histopathology with hematoxylin and eosin staining of sections. On microscopic exam, host response was categorized as to reversible or non-reversible changes. Reversible cellular changes included atrophy, depletion of zooxanthellae from gastrodermis, wound repair, hyperplasia of basal body wall, and inflammation whereas irreversible changes comprised necrosis and fragmentation. Visible organisms associated with lesions were classified as fungi, bacteria, cyanobacteria, sponges, or algae [30]. Tissue-loss lesions not found associated with obvious micro-organisms were termed ‘white syndrome’ indicating a tissue loss disease of unknown etiology. In addition, all samples collected during surveys for histology were also screened for cell-associated microbial aggregates (CAMA). Certain coral genera in the Indo-Pacific contain CAMAs that are proposed to be facultative secondary symbionts important in coral health [31] and so this was an opportunity to examine whether Red Sea corals also contained CAMAs.

### Statistical analyses

Underwater time constraints prevented enumeration of all coral colonies within the wider belt transects surveyed for disease. Hence, prevalence of lesions was determined by extrapolating colony counts within the 25 x 1 m transect to the wider 25 x 6 m disease survey area and by using this as the denominator of prevalence calculations (e.g. (number of colonies with lesions/ total number of estimated colonies) * 100). Overall prevalence was the percentage of colonies surveyed that had a particular lesion type with all surveys combined. The frequency of disease occurrence (FOC) reflects the spatial distribution of diseases on reefs and was defined as the number of sites having corals with lesions divided by total number of sites surveyed. The denominator for FOC calculations were limited to sites that had the specific coral taxa exhibiting lesions. All calculations for disease prevalence or FOC were by coral genera (e.g. prevalence of *Porites* growth anomalies was calculated as the total number of affected *Porites* colonies divided by the total number of *Porites* colonies surveyed, multiplied by 100. Percent bleaching was calculated as the number of points with bleached cover divided by the total number of points (bleached +healthy). Climatology data for each survey site were obtained from NOAA’s Coral Reef Watch Product Suite Version 3.1 [32,33] and include the average minimum and maximum sea surface temperatures (SSTs) over the last 25 years (historical SSTs), and the degree heating weeks (DHW) for the 12 week period prior to Oct. 1, 2015.

Data were not normally distributed, even with transformation, so non-parametric analyses were used. A Kruskal-Wallis test and Dunn’s post hoc tests were used to examine regional (Al Lith, Thuwal, Yanbu) differences in coral cover, number of coral genera, colony densities, disease prevalence, percent bleaching and degree heating weeks (DHW). Analysis of similarities (ANOSIM) were performed (999 permutations) on weighted (presence and abundance of coral taxa) Bray-Curtis similarity matrices to test for significant differences in coral communities between regions (Al Lith, Thuwal, Yanbu) using PRIMER-E v7 (Primer-E Ltd.). Weighted nMDS plots based on Bray-Curtis similarity matrices were produced to visualize regional differences. Disease susceptibility among coral taxa was examined using a chi-square test for equality of distributions comparing the distribution of the number of diseased versus healthy colonies among the coral genera affected by disease. The relationship between disease prevalence and three potential co-factors: coral cover, percent bleaching and DHW, was examined using a Spearman’s rank order correlation. Non-parametric statistics were performed using JMP Pro 13 statistical software (SAS Institute Inc., Buckinghamshire, UK). The map indicating survey locations was created using reefMapMaker [34].

## Results

### Coral reef characteristics and coral community structure

For the 22 sites surveyed, overall average coral cover was 44.7% (range 3-83%), average soft coral cover was 13.7% (range 0-45%), average crustose coralline algae (CCA) cover was 7.2% (range 0-30%), and average macroalgae cover was <1%. Average colony density was 16.2/m^2^ (range 7.3-28.2). Across all sites, 30 coral genera were identified with the three dominant coral taxa being *Porites* (20.6% of the community), *Pocillopora* (14.9%) and *Favites/Dipsastraea* (9.5%) (S2 Table).

### Regional differences in coral cover, colony densities and genera richness

There were significant differences among the three regions (Al Lith, Thuwal, Yanbu) in coral cover (Kruskal-Wallis, X^2^=11.0, df=2, p=0.004), coral colony densities (Kruskal-Wallis, X^2^=7.2, df=2, p=0.03) and number of coral genera (Kruskal-Wallis, X^2^=13.1, df=2, p=0.001) (Fig 2). Coral communities were also significantly different among regions (ANOSIM, Global R = 0.13, p = 0.047) (Fig 3). Coral communities in Al Lith differed significantly from Yanbu (ANOSIM, Global R = 0.415, p = 0.013) but Thuwal was not significantly different from Yanbu (ANOSIM, Global R = 0.046, p = 0.301) or Al Lith (ANOSIM, Global R = 0.097, p = 0.169).

**Fig 2.**
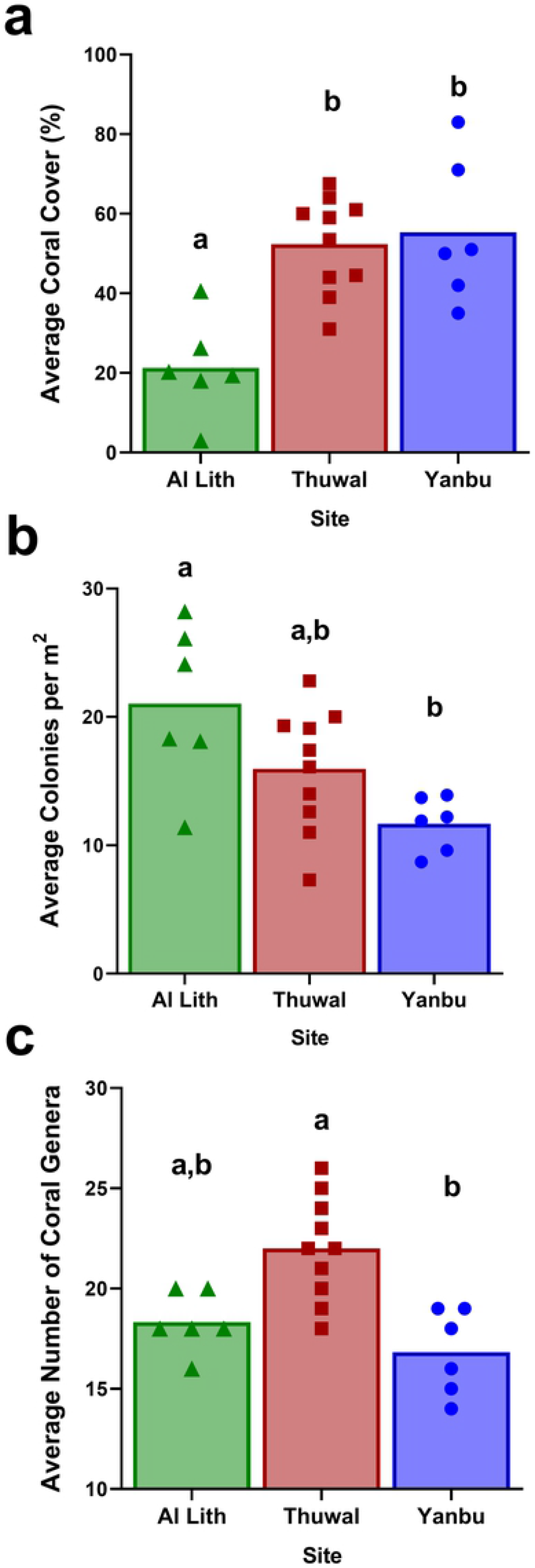
Regional differences in coral cover, colony densities and coral genus richness. Letters indicate results of Dunn’s multiple group comparison tests. Six sites each were surveyed in Al Lith and Yanbu and ten sites in Thuwal.

**Fig 3.**
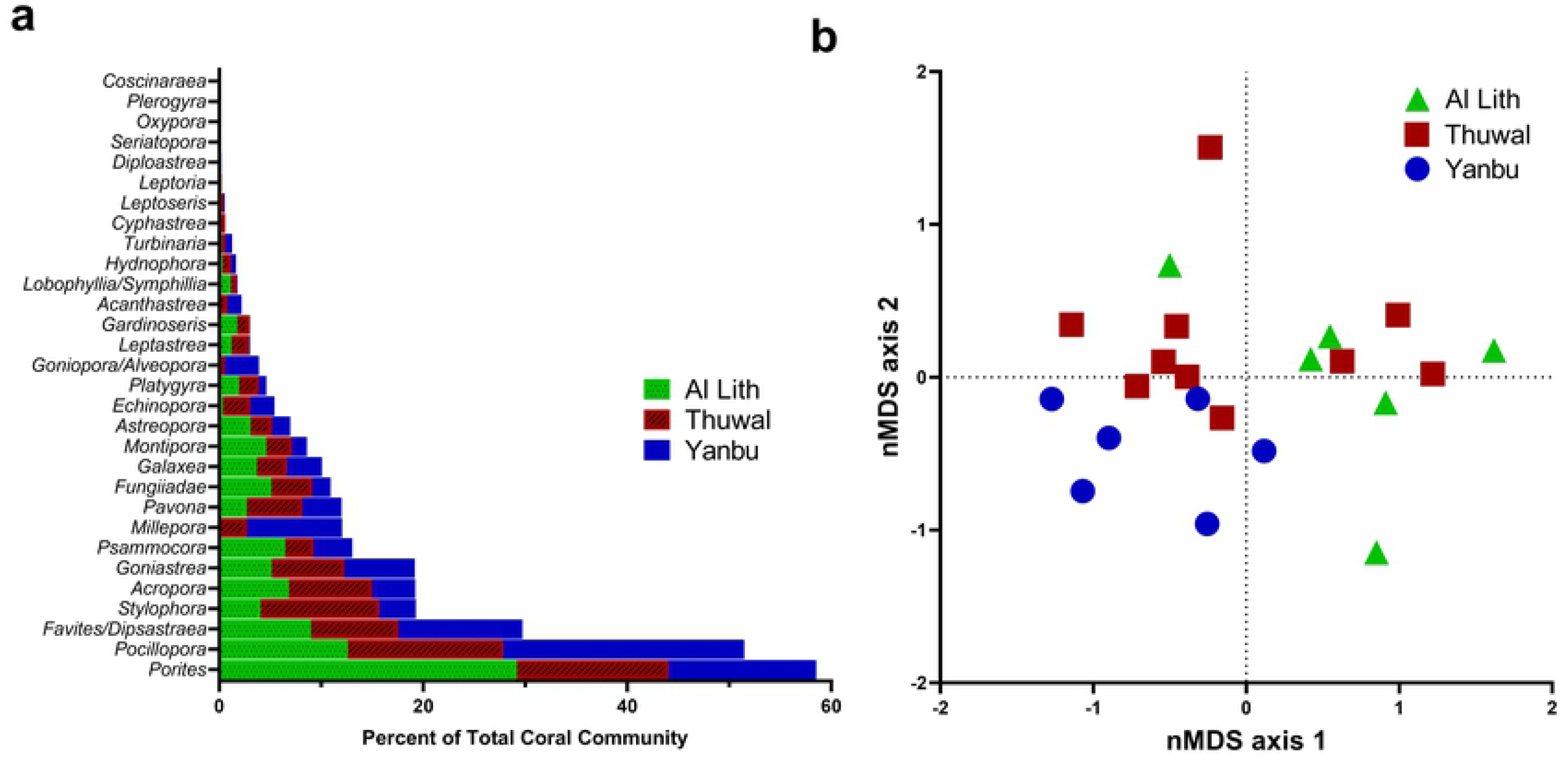
A) Average relative abundance of coral taxa in three regions surveyed for coral disease in 2015. Data show the average percent of the coral community represented by each coral genus. B) A non-metric multi-dimensional scaling (nMDS plot) illustrating the differences in coral communities between regions. Six sites each were surveyed in Al Lith and Yanbu and ten sites in Thuwal.

### Regional differences in percent bleaching and amount of heat stress

There were significant differences among the three regions (Al Lith, Thuwal, Yanbu) in percent bleaching (Kruskal-Wallis, X^2^=14.0, df=2, p=0.0009) (Table 1). Sites in the Al Lith region had the highest proportion of bleached coral cover (avg. 33.5%) followed by sites in Thuwal (avg. 13.1%) with no bleaching found at sites in Yanbu. There were also significant differences among the three regions in DHW (Kruskal-Wallis, X^2^=13.4, df=2, p=0.001)(Table 1). Sites in Yanbu had the highest DHW (avg. 4.9), followed by Al Lith (avg. 4.4) and Thuwal (avg. 3.6). No relationship was found between percent bleaching and DHW (Spearman’s rank, Pho=-0.24, p=0.3).

**Table 1.**
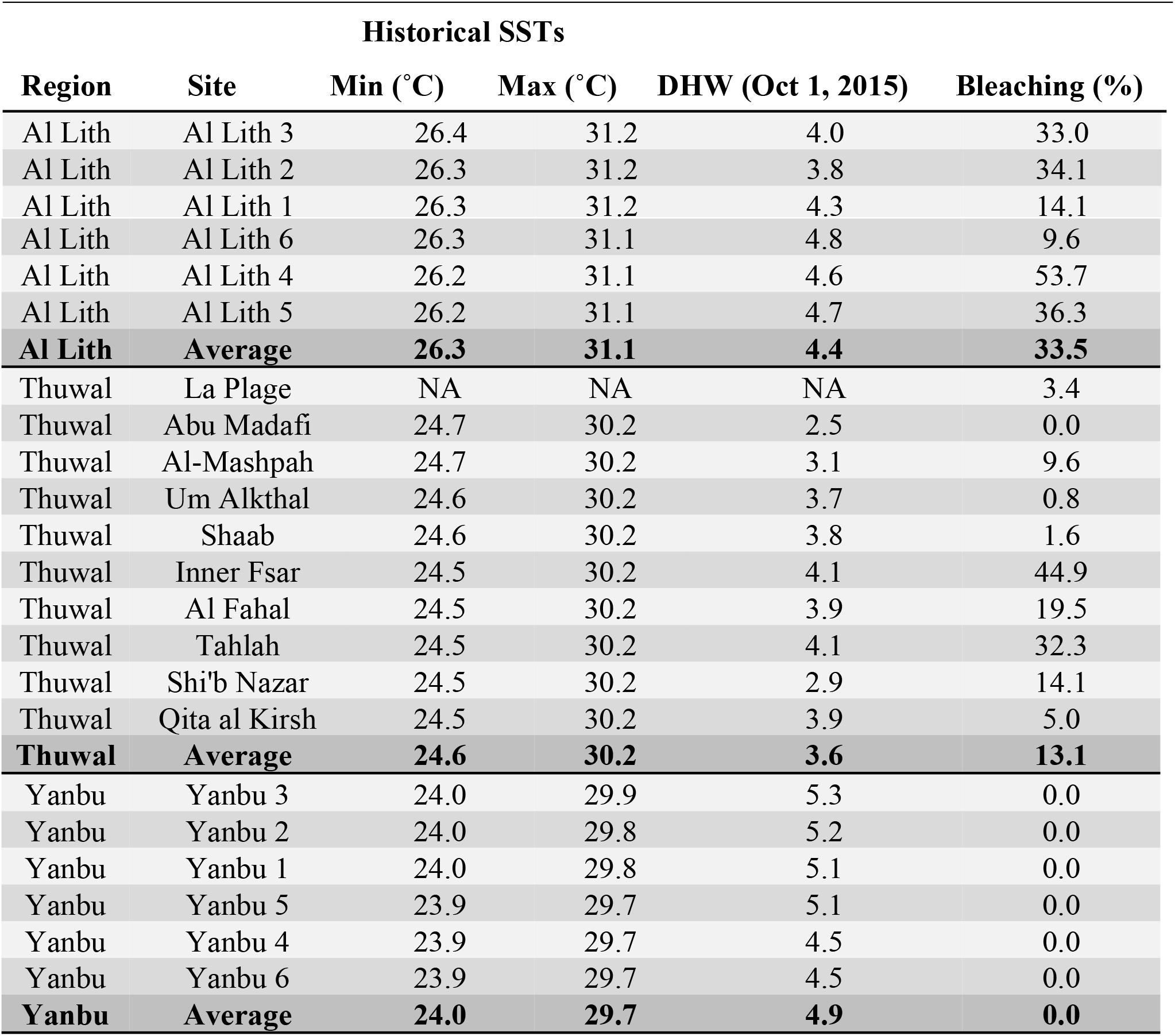
Summary of temperature variability and bleaching response in different study sites along the Saudi coast of the Red Sea. Avg. minimum and maximum SSTs over the last 25 years, the Degree Heating Weeks (DHW) for the 12-week period prior to Oct. 1, 2015, and degree of bleaching is reported for each survey site.

| Region | Site | Historical SSTs |  |  |  |
| --- | --- | --- | --- | --- | --- |
|  |  | Min (°C) | Max (°C) | DHW (Oct 1, 2015) | Bleaching (%) |
| Al Lith | Al Lith 3 | 26.4 | 31.2 | 4.0 | 33.0 |
| Al Lith | Al Lith 2 | 26.3 | 31.2 | 3.8 | 34.1 |
| Al Lith | Al Lith 1 | 26.3 | 31.2 | 4.3 | 14.1 |
| Al Lith | Al Lith 6 | 26.3 | 31.1 | 4.8 | 9.6 |
| Al Lith | Al Lith 4 | 26.2 | 31.1 | 4.6 | 53.7 |
| Al Lith | Al Lith 5 | 26.2 | 31.1 | 4.7 | 36.3 |
| **Al Lith** | **Average** | **26.3** | **31.1** | **4.4** | **33.5** |
| Thuwal | La Plage | NA | NA | NA | 3.4 |
| Thuwal | Abu Madafi | 24.7 | 30.2 | 2.5 | 0.0 |
| Thuwal | Al-Mashpah | 24.7 | 30.2 | 3.1 | 9.6 |
| Thuwal | Um Alkthal | 24.6 | 30.2 | 3.7 | 0.8 |
| Thuwal | Shaab | 24.6 | 30.2 | 3.8 | 1.6 |
| Thuwal | Inner Fsar | 24.5 | 30.2 | 4.1 | 44.9 |
| Thuwal | Al Fahal | 24.5 | 30.2 | 3.9 | 19.5 |
| Thuwal | Tahlah | 24.5 | 30.2 | 4.1 | 32.3 |
| Thuwal | Shi'b Nazar | 24.5 | 30.2 | 2.9 | 14.1 |
| Thuwal | Qita al Kirsh | 24.5 | 30.2 | 3.9 | 5.0 |
| **Thuwal** | **Average** | **24.6** | **30.2** | **3.6** | **13.1** |
| Yanbu | Yanbu 3 | 24.0 | 29.9 | 5.3 | 0.0 |
| Yanbu | Yanbu 2 | 24.0 | 29.8 | 5.2 | 0.0 |
| Yanbu | Yanbu 1 | 24.0 | 29.8 | 5.1 | 0.0 |
| Yanbu | Yanbu 5 | 23.9 | 29.7 | 5.1 | 0.0 |
| Yanbu | Yanbu 4 | 23.9 | 29.7 | 4.5 | 0.0 |
| Yanbu | Yanbu 6 | 23.9 | 29.7 | 4.5 | 0.0 |
| **Yanbu** | **Average** | **24.0** | **29.7** | **4.9** | **0.0** |

### Disease prevalence, frequency of occurrence and types of diseases

An estimated 75,750 coral colonies were examined for disease and the overall disease prevalence (all sites combined) was 0.17%. A total of 21 diseases in 16 coral taxa were recorded (Table 2). Lesion types included tissue loss diseases of unknown etiology (white syndromes), growth anomalies, distinct focal bleached patches, skeletal eroding band (folliculinid ciliate disease), black band disease (tissue loss due to microbial consortium dominated by filamentous cyanobacteria) and endolithic hypermycosis (purple discoloration due to endolithic fungal infection) (Fig 4). The three most common diseases were *Acropora* white syndrome found at 59.1% of the survey sites, *Porites* growth anomalies found at 40.9% of the sites, and *Porites* white syndrome found at 31.8% of the sites.

**Fig 4.**
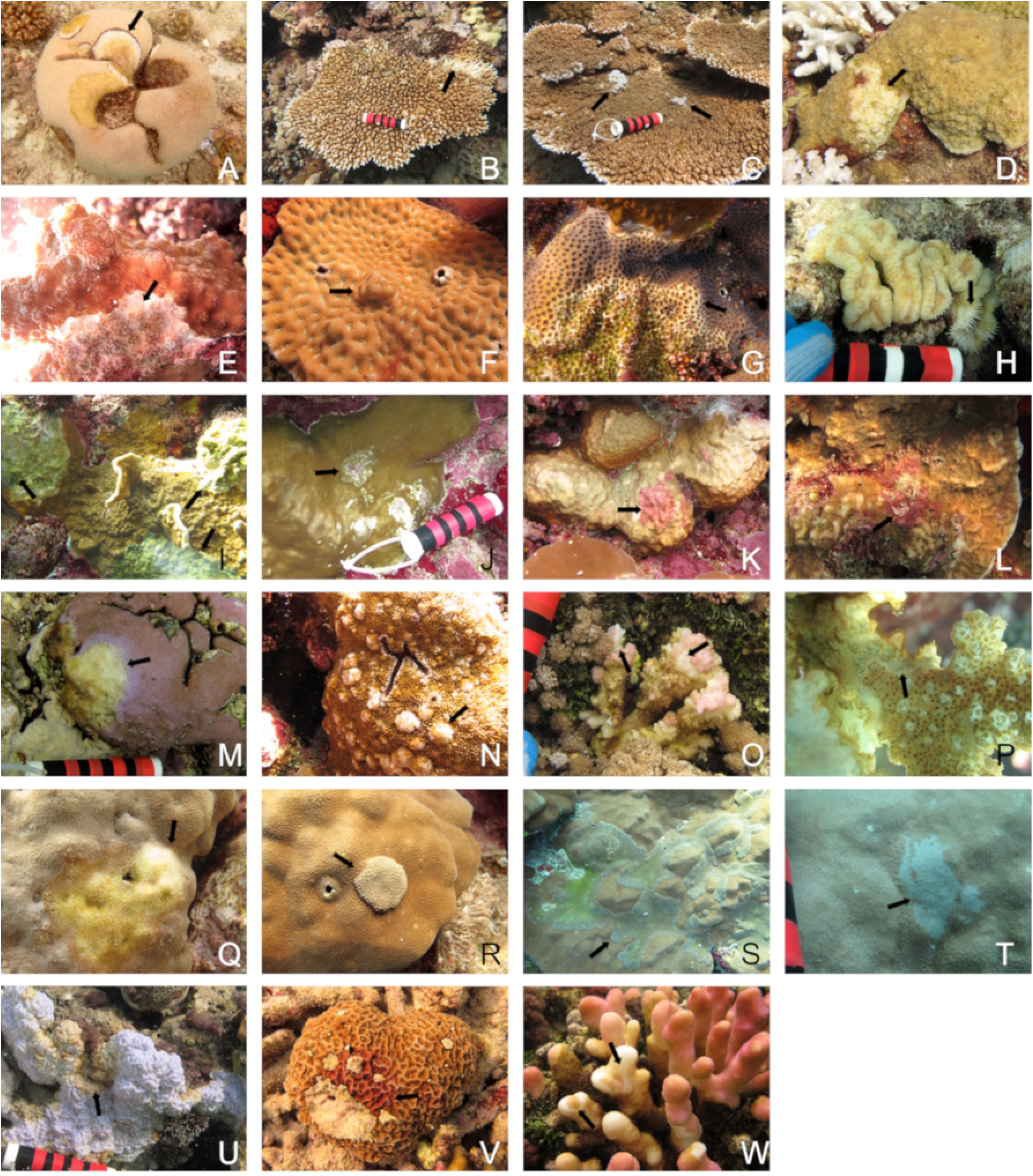
Examples of different coral diseases encountered during disease surveys along the Saudi coast of the Red Sea. WS=white syndrome, GA=growth anomaly, EH=endolithic hypermycosis, Bl=bleached. A) black band disease, B) *Acropora* WS, C) *Acropora* GA, D) *Astreopora* WS, E) *Echinopora* WS, F) *Favites* GA, G) *Goniastrea* WS, H) *Lobophyllia* WS, I) *Millepora* WS, J) *Millepora* GA, K) *Millepora* EH, L) *Pavona* EH, M) *Montipora* WS, N) *Montipora* GA, O) *Pocillopora* WS, P) *Pocillopora* SEB, Q) *Porites* WS, R) *Porites* GA, S) *Porites* GA, T) *Porites* Bl patch, U) *Psammocora* WS, V) *Psammocora* EH, W) *Stylophora* WS

**Table 2.**
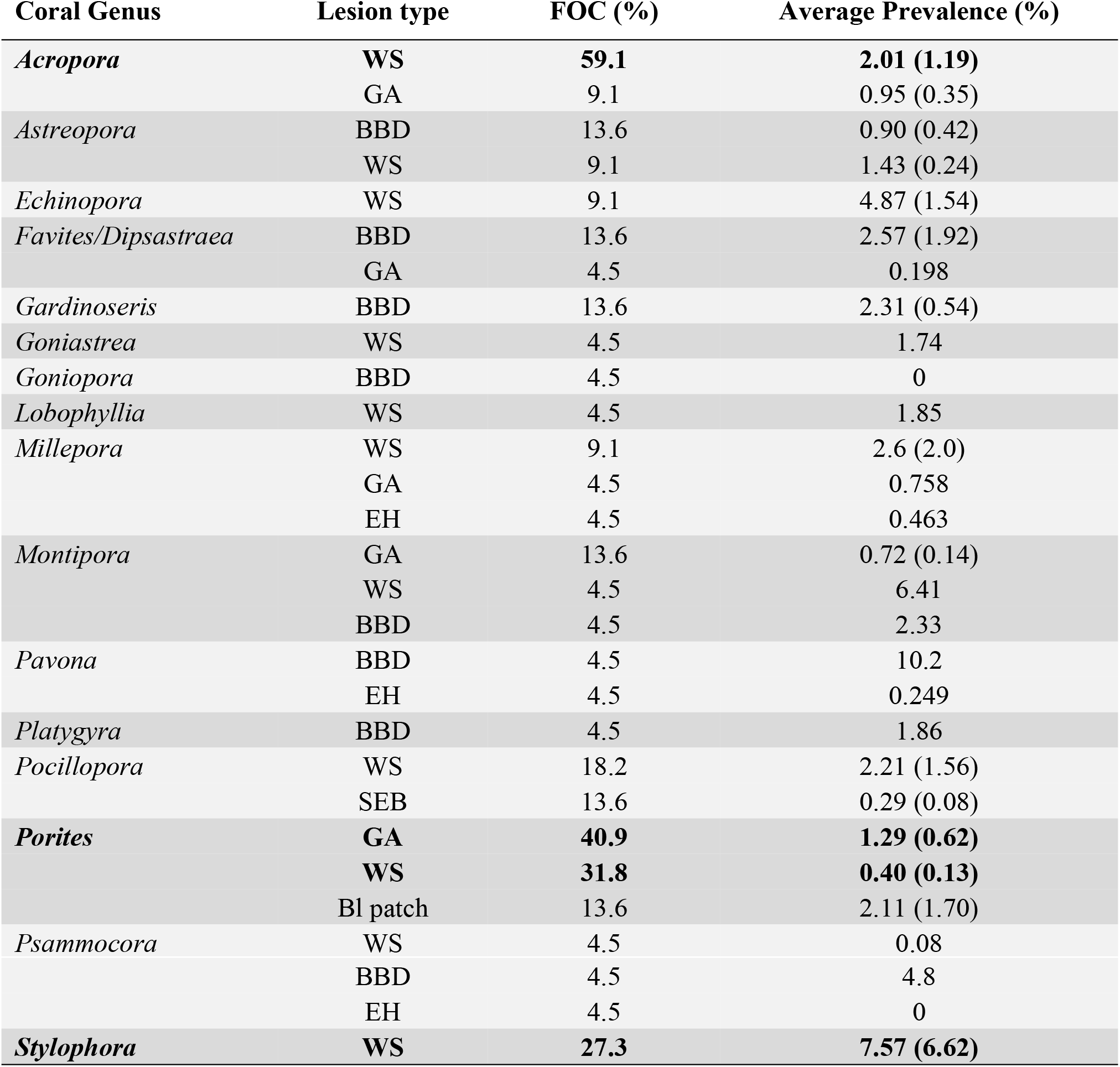
Frequency of occurrence (FOC) and average prevalence of coral diseases found during surveys. FOC represents the proportion of sites containing corals with each respective disease. Average prevalence (standard error in parentheses) calculated exclusively from the sites containing each respective disease. Absence of a standard error indicates the disease was found at a single survey site. Prevalence data includes diseased colonies only within transects and so will differ from frequency of disease occurrence data. WS=white syndrome, GA=growth anomaly, SEB=skeletal eroding band (ciliates), BBD=black band disease, Bl patch=focal bleached area, EH=endolithic hypermycosis. The most common diseases are in bold.

| Coral Genus | Lesion type | FOC (%) | Average Prevalence (%) |
| --- | --- | --- | --- |
| *Acropora* | **WS** | **59.1** | **2.01 (1.19)** |
|  | GA | 9.1 | 0.95 (0.35) |
| *Astreopora* | BBD | 13.6 | 0.90 (0.42) |
|  | WS | 9.1 | 1.43 (0.24) |
| *Echinopora* | WS | 9.1 | 4.87 (1.54) |
| *Favites/Dipsastraea* | BBD | 13.6 | 2.57 (1.92) |
|  | GA | 4.5 | 0.198 |
| *Gardinoseris* | BBD | 13.6 | 2.31 (0.54) |
| *Goniastrea* | WS | 4.5 | 1.74 |
| *Goniopora* | BBD | 4.5 | 0 |
| *Lobophyllia* | WS | 4.5 | 1.85 |
| *Millepora* | WS | 9.1 | 2.6 (2.0) |
|  | GA | 4.5 | 0.758 |
|  | EH | 4.5 | 0.463 |
| *Montipora* | GA | 13.6 | 0.72 (0.14) |
|  | WS | 4.5 | 6.41 |
|  | BBD | 4.5 | 2.33 |
| *Pavona* | BBD | 4.5 | 10.2 |
|  | EH | 4.5 | 0.249 |
| *Platygyra* | BBD | 4.5 | 1.86 |
| *Pocillopora* | WS | 18.2 | 2.21 (1.56) |
|  | SEB | 13.6 | 0.29 (0.08) |
| *Porites* | **GA** | **40.9** | **1.29 (0.62)** |
|  | **WS** | **31.8** | **0.40 (0.13)** |
|  | Bl patch | 13.6 | 2.11 (1.70) |
| *Psammocora* | WS | 4.5 | 0.08 |
|  | BBD | 4.5 | 4.8 |
|  | EH | 4.5 | 0 |
| *Stylophora* | WS | 27.3 | 7.57 (6.62) |

### Histopathology of coral lesions

Paired normal and abnormal tissues collected from 43 colonies representing 15 coral genera were examined. This included samples from 13 *Porites* colonies (30% of the samples), 6 *Stylophora* (14%), 5 *Pocillopora* (12%), 3 each from *Acropora*, *Astreopora*, and *Psammocora* (7% each), 2 *Gardinoseris* (5%), and 1 each from *Dipsastraea, Echinopora, Favites, Goniastrea, Goniopora, Leptoseris*, *Montipora*, and *Sarcophyton* (2% each). The most common lesion sampled for histology was tissue loss (67%) (Figs 5A-C), followed by growth anomalies (16.5%) (Figs 5D-E) and discoloration (16.5%) (Fig 5F). Of the 29 colonies with tissue loss, the tissue loss lesions could be further subdivided as subacute (n=14) (Fig 5A), acute (n=10) (Fig 5B), chronic (n=3) (Fig 5C), or a combination of the three (n=2). Histology samples originated from Yanbu (n=13), Al Lith (n=11), and Thuwal (n=19).

**Fig 5.**
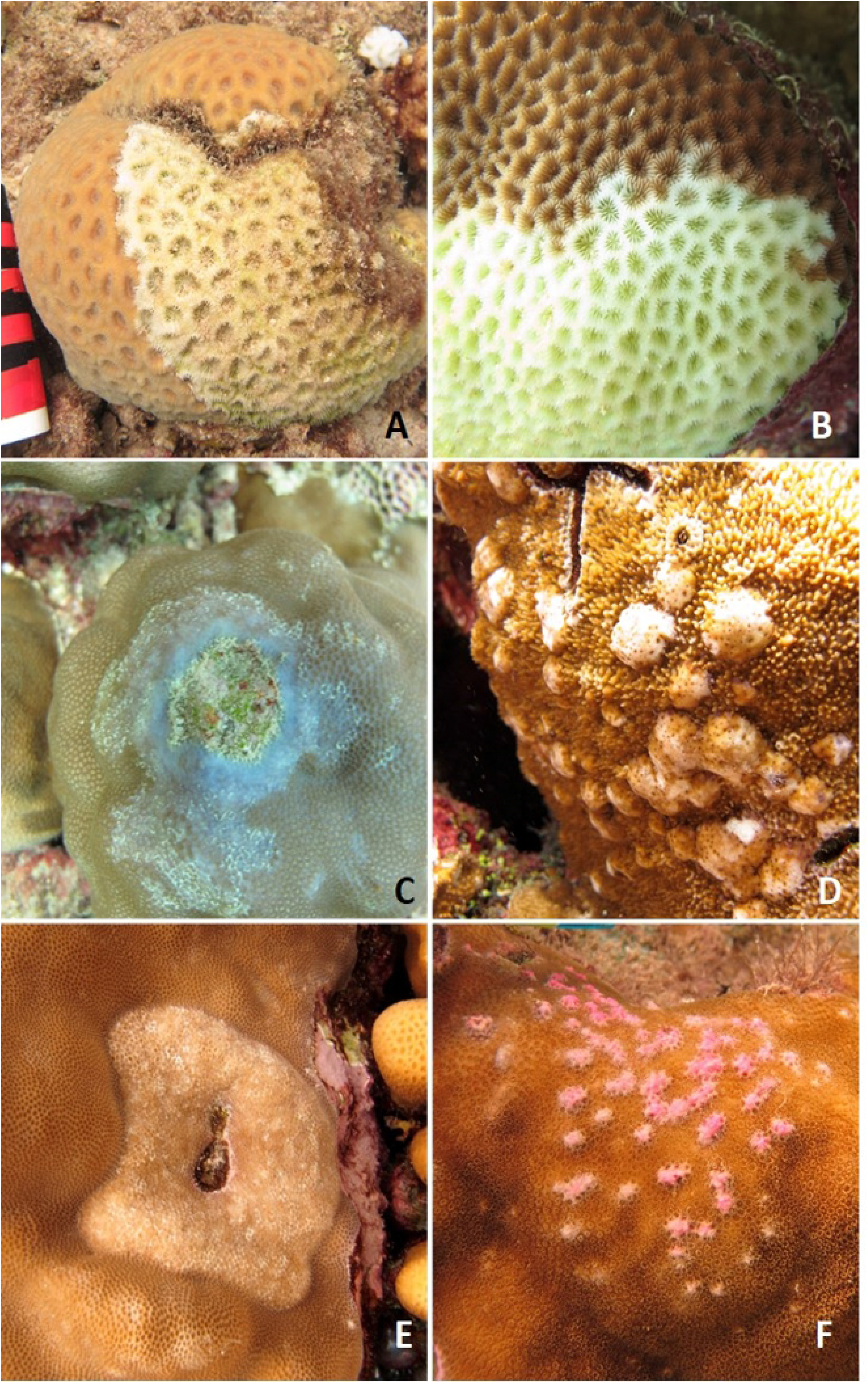
Representative types of lesions sampled for histopathology. A) Acute tissue loss, *Dipsastraea* sp., B) Acute tissue loss, *Goniastrea* sp., C) Chronic tissue loss with discoloration, *Porites* sp., D) Growth anomaly, *Montipora* sp., E) Growth anomaly, *Porites* sp., F) Discoloration, *Porites* sp.

A total of 87 tissue samples were collected from colonies manifesting tissue loss (N=29), growth anomalies (n=9) and discoloration (n=7) with the remainder being from grossly apparently normal tissues. Of 29 histology sections from colonies with tissue loss, 12 had necrosis either alone or associated with fungi, cyanobacteria, algae, or sponges, 6 sections had atrophy of tissues with depletion of zooxanthellae, 5 had no evident microscopic changes, and 3 had varying degrees of inflammation associated with algae, fungi, or cyanobacteria. For 9 samples from growth anomalies, 5 had no evident changes, 3 had hyperplasia of basal body wall, and 1 had necrosis with algae. Of 7 sections with discoloration, all but 2 had necrosis with inflammation, fungi, or algae with the 2 remaining with no evident lesions. Of 42 apparently normal fragments, 16 had no evident changes, 12 had necrosis associated with fungi, cyanobacteria, algae or sponges, 8 had atrophy and depletion of zooxanthellae, 4 had inflammation sometimes associated with algae, and 1 had endolithic fungi. Of organisms associated with host response (inflammation, necrosis), fungi dominated (n=12) followed by cyanobacteria (n=6), algae (n=3), and sponges (n=1).

### Cell associated microbial aggregates (CAMA)

A total of 87 coral fragments from 15 genera were examined histologically for CAMAs. CAMAs were found in four coral genera, including 5 out of 6 of the *Stylophora* fragments (83% of the samples examined) (Fig 6A), three out of 13 *Porites* fragments (21%)(Fig 6B), two out of three *Acropora* fragments (67%)(Fig 6C) and two out of five *Pocillopora* fragments (33%)(Fig. 6D).

**Fig 6.**
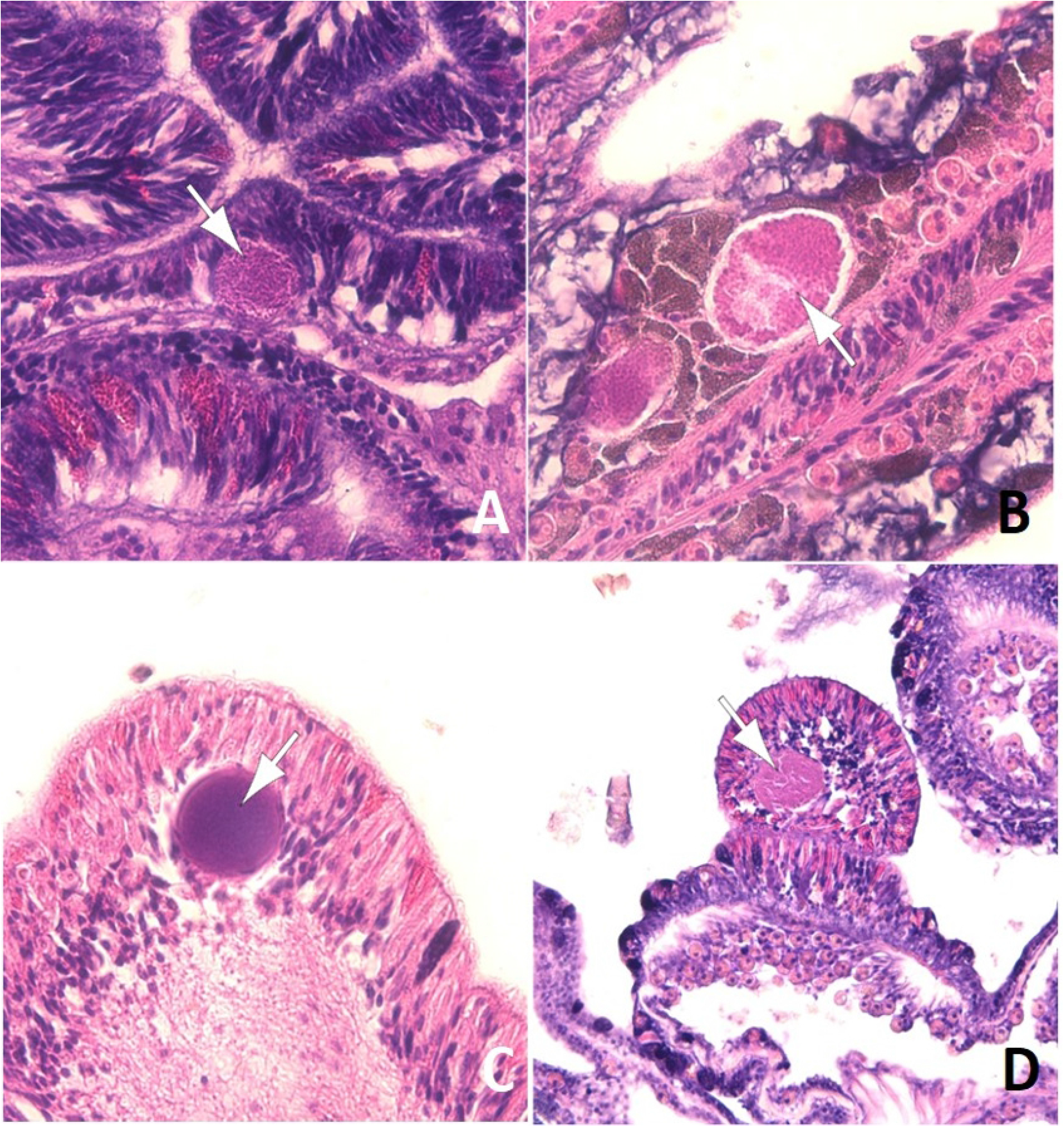
Cell associated microbial aggregates (arrows) in epidermis (A, C, D) and gastrodermis (B) of *Stylophora* (A), *Porites* (B), *Acropora* (C) and *Pocillopora* (D).

### Differences in disease prevalence among coral genera

Out of 30 coral genera found within transects, 16 had lesions indicative of disease. Disease prevalence diffed among coral genera (X^2^=90.3, df=16, p<0.005) with *Acropora* having the highest overall disease prevalence (0.54%), followed by *Millepora* (0.44%) and *Lobophyllia* (0.38%) (Fig 7).

**Fig 7.**
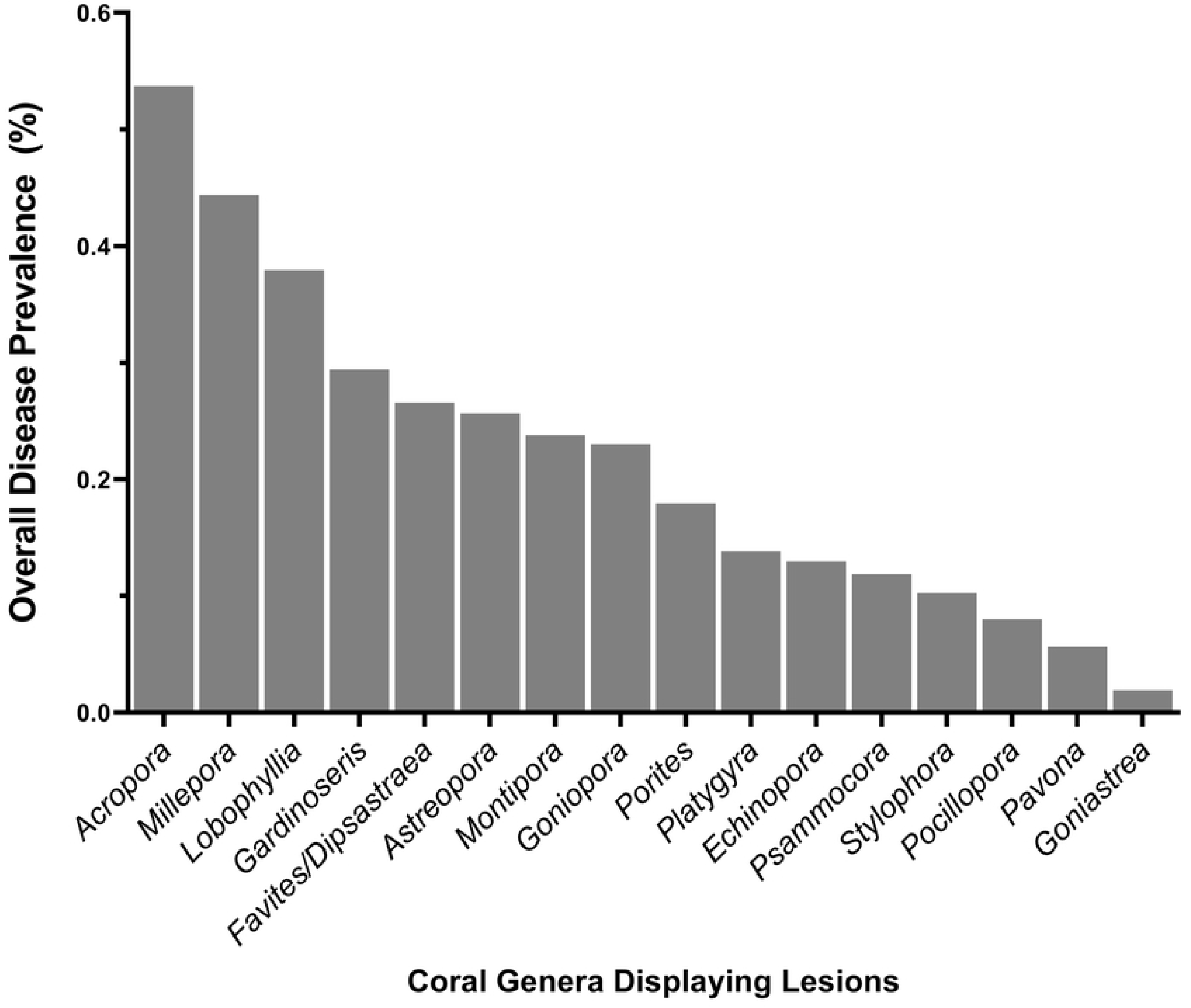
Differences in overall disease prevalence among coral taxa. Data show overall prevalence with all surveys combined.

### Differences in disease among regions

Average disease prevalence differed significantly among regions (Kruskal-Wallis, X^2^=6.6, df=2, p=0.04) (Fig 8) with differences among specific diseases in the frequency of occurrence and average prevalence (Table 3). Of the three most common lesions (BBD, WS, growth anomalies), there were significant regional differences in black band disease (Kruskal-Wallis, X^2^=6.3, df=2, p=0.04), and white syndrome (Kruskal-Wallis, X^2^=10.2, df=2, p=0.006) but not growth anomalies (Kruskal-Wallis, X^2^=4.6, df=2, p=0.1). Average BBD prevalence was highest in Al Lith (0.33% SE±0.3), although mainly due to one outbreak site, followed by Thuwal (0.009% SE±0.005), and 0% in Yanbu. Average white syndrome prevalence was highest in Yanbu (0.47% SE±0.15) followed by Al Lith (0.06% SE±0.02) and Thuwal (0.07% SE±0.03). Average prevalence of growth anomalies was 0.11% (SE±0.04) in Yanbu, 0.04% (SE±0.02) in Thuwal and 0.003% (SE±0.003) in Al Lith.

**Fig 8.**
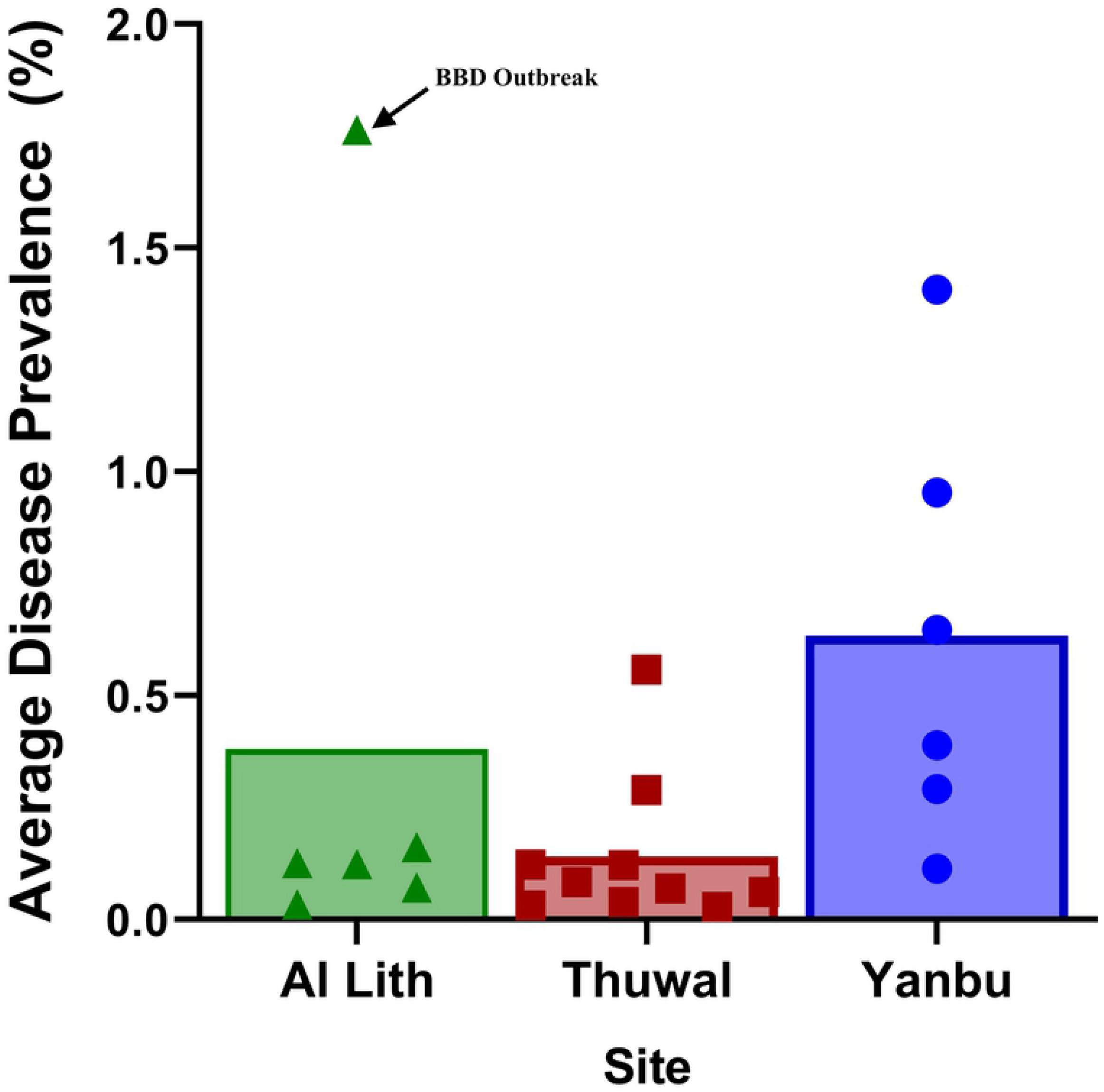
Regional differences in disease prevalence at sites surveyed along the Saudi Arabian coast of the Red Sea in October-November 2015. Letters indicate results of Dunn’s multiple group comparison tests. Six sites each were surveyed in Al Lith and Yanbu and ten sites in Thuwal.

**Table 3.** Regional differences in frequency of occurrence (FOC) and average prevalence (±SEM) of different coral diseases found during surveys in the central Red Sea. Prevalence data includes diseased colonies only within transects and so will differ from frequency of disease occurrence data. WS=white syndrome, GA=growth anomaly, SEB=skeletal eroding band (ciliates), BBD=black band disease, Bl patch=focal bleached area, EH=endolithic hypermycosis. Six sites each were surveyed in Al Lith and Yanbu and ten sites in Thuwal.

| Coral Genus | Lesion Type | FOC (%) |  |  | Average Disease Prevalence (%) |  |  |
| --- | --- | --- | --- | --- | --- | --- | --- |
|  |  | Yanbu | Thuwal | Al Lith | Yanbu | Thuwal | Al Lith |
| *Montipora* | BBD | 0 | 0 | 16.7 | 0 | 0 | 2.33 |
| *Favites/Dipsastraea* | BBD | 0 | 10 | 33.3 | 0 | 0.5 | 3.6 (2.8) |
| *Psammocora* | BBD | 0 | 0 | 16.7 | 0 | 0 | 4.8 |
| *Gardinoseris* | BBD | 0 | 10 | 33 | 0 | 3.3 | 1.81 (0.29) |
| *Astreopora* | BBD | 0 | 10 | 33.3 | 0 | 0.32 | 1.19 (0.52) |
| *Pavona* | BBD | 0 | 0 | 16.7 | 0 | 0 | 10.2 |
| *Platygyra* | BBD | 0 | 0 | 16.7 | 0 | 0 | 1.9 |
| *Goniopora* | BBD | 0 | 0 | 16.7 | 0 | 0 | 0 |
| *Porites* | Bl patch | 16.7 | 20 | 0 | 5.1 | 0.42 (0.19) | 0 |
| *Millepora* | EH | 0 | 10 | 0 | 0 | 0.8 | 0 |
| *Psammocora* | EH | 0 | 10 | 0 | 0 | 0 | 0 |
| *Pavona* | EH | 0 | 10 | 0 | 0 | 0.25 | 0 |
| *Acropora* | GA | 0 | 20 | 0 | 0 | 0.95 (0.35) | 0 |
| *Montipora* | GA | 0 | 10 | 0 | 0 | 0.72 (0.14) | 0 |
| *Porites* | GA | 66.7 | 40 | 0 | 2.15 (1.13) | 0.56 (0.23) | 0 |
| *Millepora* | GA | 0 | 10 | 0 | 0 | 0.8 | 0 |
| *Favites/Dipsastraea* | GA | 0 | 10 | 0 | 0 | 0.2 | 0 |
| *Pocillopora* | SEB | 33.3 | 10 | 0 | 0.33 (0.12) | 0.2 | 0 |
| *Acropora* | WS | 33.3 | 80 | 50 | 8.4 (7.6) | 0.98 (0.42) | 0.5 (0.23) |
| *Pocillopora* | WS | 50 | 0 | 16.7 | 0.87 (0.55) | 0 | 8.3 |
| *Montipora* | WS | 33.3 | 0 | 0 | 6.41 | 0 | 0 |
| *Porites* | WS | 50 | 20 | 16.7 | 0.67 (0.04) | 0.04 (0.04) | 0.15 (0.03) |
| *Echinopora* | WS | 33.3 | 0 | 0 | 4.86 (1.56) | 0 | 0 |
| *Millepora* | WS | 33.3 | 0 | 0 | 2.6 (2.0) | 0 | 0 |
| *Goniastrea* | WS | 16.7 | 0 | 0 | 1.74 | 0 | 0 |
| *Psammocora* | WS | 0 | 0 | 16.7 | 0 | 0 | 0.08 |
| *Lobophyllia* | WS | 0 | 0 | 16.7 | 0 | 0 | 1.85 |
| *Astreopora* | WS | 0 | 10 | 16.7 | 0 | 0 | 1.67 |
| *Stylophora* | WS | 16.7 | 10 | 0 | 1.74 (0.74) | 0.19 (0.16) | 0.08 |

### Relationship between disease prevalence and environmental factors

We examined the relationship between disease prevalence and coral cover, percent bleaching and degree heating weeks (DHW). No significant relationship was found between disease prevalence and coral cover (Spearman’s rank, Pho= −0.06, p=0.79) or DHW (Spearman’s rank, Pho=0.35, p=0.12). A negative relationship was found between disease prevalence and coral bleaching (Spearman’s rank, Pho= −0.51, p=0.02)(Fig 9).

**Fig 9.**
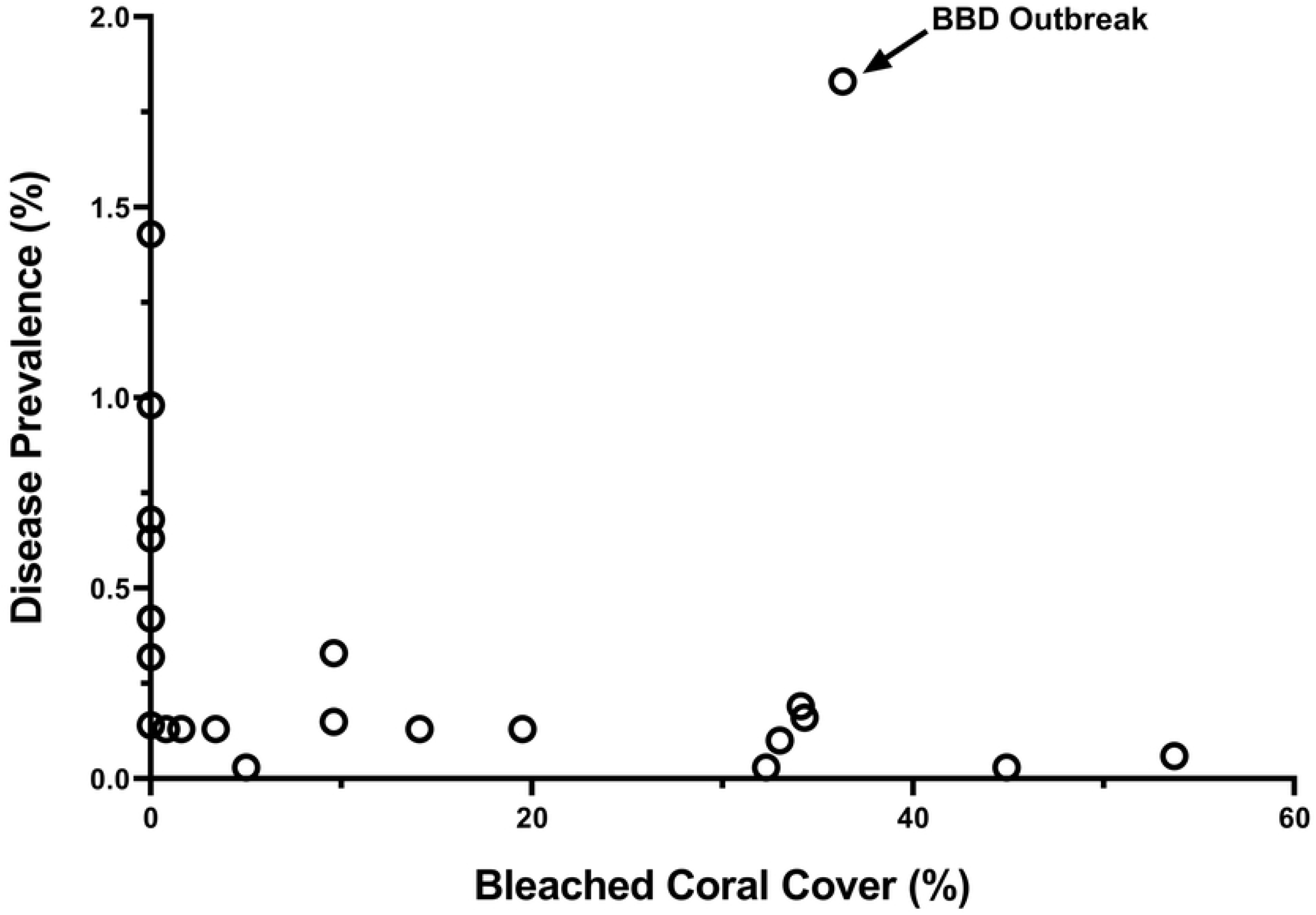
Relationship between coral disease and level of bleaching for 22 sites surveyed in 2015 along the Red Sea coast of Saudi Arabia.

## Discussion

Coral reefs are in decline globally and disease has played a significant factor in that decline [1,35,36]. Comparatively little research has been done on coral disease in the Red Sea and our study presents important information on types of diseases present on coral reefs along the Saudi Arabian Red Sea coast, prevalence of diseases, susceptible coral taxa within this region and a description of the histology of different coral lesions. Baseline data is particularly relevant considering the planned mega-building projects such as NEOM (https://www.neom.com) and the Red Sea project (https://visiontoreality.theredsea.sa), which are expected to exert heavy impacts on surrounding coral reef ecosystems. Twenty-two reefs were surveyed in the Red Sea along the Saudi Arabian coast, and robust hard coral (45%) and soft coral (14%) cover, and very low levels of macroalgae cover (<1%) were found. Thirty hard coral genera were found within transects. Coral reefs had widespread but overall low prevalence of disease (<0.5%) with 20 diseases recorded affecting 16 coral taxa and disease lesions found on corals at all sites surveyed. Coral reefs in the Red Sea had diseases typical of many regions including black band disease, white syndromes, endolithic hypermycosis, skeletal eroding band, growth anomalies and distinct focal bleached patches.

### Overview of diseases affecting corals in the Red Sea

#### Black band disease (BBD)

BBD has been reported from coral reefs across the world [37] including the Red Sea [38,39]. BBD typically remains at low background levels [40,41] with seasonal outbreaks occurring [42–44]. At our sites in the Red Sea, there was a similar pattern with a low prevalence of infected corals (0.02-0.12%) found at seven out of 22 sites. Antonius [45] found similar levels of BBD along the Saudi Arabian coast in the 1980s suggesting BBD levels have not changed significantly along these reefs in the past 30 years. We also documented outbreak levels at one site in the Al Lith region, which has been described elsewhere [38]. In fact, most of the BBD cases were found at sites in Al Lith whereas no cases were found in the northern-most region (Yanbu). BBD is sensitive to water temperatures becoming more common in summer months when water temperature and light levels are higher and usually disappears during colder winter months [46–48]. BBD infections also appear following bleaching events[49,50]. Al Lith has higher average SST ranges (26.3°C to 31.1°C) as compared to Yanbu with average SSTs ranging from 24.0°C to 29.7°C. Al Lith also had the highest level of bleaching among the regions surveyed whereas no bleaching was found in Yanbu. Thus, higher average SSTs combined with a higher bleaching potential may leave reefs at Al Lith particularly vulnerable to BBD infections.

#### White syndromes (WS)

Tissue loss diseases of unknown etiology (white syndromes) are commonly found on a multitude of species on reefs throughout the world [51] and white syndromes were found in all three regions affecting 10 coral genera. *Acropora* white syndrome was the most widespread disease occurring in all regions at 13 of the 17 reefs surveyed. In addition, *Acropora* white syndrome occurred at outbreak levels at three sites with the highest level at Yanbu 2 (16% prevalence), followed by Abu Madafi (3.6% prevalence) and Al-Mashpah (1.9% prevalence) (both Thuwal region). We defined a site as having a localized disease outbreak if prevalence was higher than the overall prevalence for this region which was <1%. *Acropora* is an exceptionally vulnerable coral genus to tissue loss diseases throughout the world [51] and the Red Sea is no exception.

#### Endolithic hypermycosis (EH)

Endolithic hypermycosis was an uncommon disease with only three cases noted within three different coral genera (*Millepora, Psammocora, Pavona*). In other regions, this lesion is associated with overgrowth of coral tissue by endolithic fungi [52–54] and the samples we examined also had a consistent histological diagnosis of endolithic fungal invasion. This disease has been reported from American Samoa [55], Hawaii [52], Micronesia [54] and New Caledonia [53] and so the present study extends this disease to the reefs of the Red Sea (biogeographic range extension). There are no prior reports of EH in *Millepora* and so the present report also potentially extends the affected taxa to include *Millepora,* a skeleton-forming hydrozoan. However, although the gross lesion on *Millepora* was consistent with endolithic hypermycosis, histology was not done; therefore, confirming presence of the fungus requires future microscopic examination.

#### Skeletal eroding band (SEB)

Skeletal eroding band (SEB) was found exclusively on pocilloporids. SEB is caused by folliculinid ciliates with tissue loss occurring when motile larval stages migrate into the tissue edges, and secrete pseudochitinous loricae which embed in the coral skeleton [56]. The disease is characterized by a dark band of varied width, adjacent to the healthy tissue, with the denuded skeleton behind the band littered with discarded black loricae [57]. Folliculinid ciliates readily colonize recently exposed coral skeletons [58] so presence of ciliates on coral skeletons does not necessarily indicate ciliate disease. Hence, we only scored a lesion as SEB if we found ciliates within millimeters of live tissue and did not include tissue loss lesions with loricae in dead skeleton further from the lesion edge. However, as we did not follow tissue loss lesions through time, we cannot rule out that lesions that we scored as SEB, were opportunistic colonization of ciliates following tissue lost to other processes. Unfortunately, verified SEB infections have not been characterized histologically, so the role of folliculinids in contributing to gross lesions remains speculative.

SEB can be quite common on reefs affecting numerous coral taxa. For example, Page and Willis [58] found SEB at 90-100% of their survey reefs affecting at least 82 scleractinian species across the GBR. Winkler et al. [59] surveyed corals in the Gulf of Aqaba, Red Sea and found 28 coral taxa affected by SEB with an overall prevalence of 29% of the total colonies surveyed. In contrast, we only found SEB affecting pocilloporids, which are one of the most commonly affected coral genera [57] but no other coral taxa had SEB lesions. We also only found SEB infections at 3 of 22 survey sites with a prevalence of <0.5% at affected sites. The low SEB prevalence we found could be due to a more conservative approach to field diagnosis of the disease or the environmental conditions on the reefs we surveyed were not conducive to SEB infections. Transmission experiments conducted by Page and Willis [58] showed that ciliates could not colonize intact coral tissue but infections were initiated in coral with injuries. Page et al. [57] suggested that co-infection involving other pathogens and/or stress under specific environmental factors may be required for ciliates to become pathogenic.

#### Growth anomalies (GA)

Growth anomalies were found in five coral genera (*Porites, Acropora, Montipora, Millepora,* and *Favites/Dipsastraea*). Prior to our study, the only coral genus reported to be affected by GAs in the Red Sea was *Platygyra* sp. [60]. Our study now expands the biogeographic range of GAs in *Porites, Acropora, Montipora,* and *Favites/Dipsastraea* to the Red Sea. GAs in *Millepora* have not been reported elsewhere and so our study also expands the host range of GAs. In many regions of the world, *Acropora* and *Porites* have been found disproportionately affected by growth anomalies in the field [61] and consistent with this, *Porites* was the most common coral taxon affected by GAs at our sites. In contrast, acroporids were the 5^th^ most abundant coral taxa at our sites, yet *Acropora* GAs were only found at two of 22 sites. Aeby et al. [62] found that the environmental predictors of GAs differed between *Porites* and *Acropora* and so the environmental conditions in the Red Sea may be conducive to GAs in *Porites* but not *Acropora*. Alternatively, different species within a single genus can differ in their disease susceptibilities [63] so perhaps the specific *Acropora* species found in the Red Sea are less prone to growth anomalies compared to those in other regions.

#### Focal bleached patches

Discrete, focal bleached patches were found in *Porites* spp. at three sites. *Porites* bleached patches have been reported from the Persian Gulf and Oman Sea [64], New Caledonia [53], and the GBR [65]. This study now extends this disease to the Red Sea (biogeographic range extension). Bleached patches are thought to be due to a viral infection of symbiotic zooxanthellae [66,67] but little else is known about this disease.

#### Histology shows healthy tissues compromised and presence of CAMAs

Samples for histology were collected following periods of increased sea temperatures and accumulated heat stress in all three regions surveyed and bleaching occurred in two of the regions. Reflecting this stress, we found that 61% of normal fragments (no gross lesion) had some sort of microscopic lesion, mainly necrosis, half of which were associated with a microorganism or atrophy of tissues with depletion of zooxanthellae (bleaching). Indeed, the breakdown of histologic lesions for apparently normal coral fragments was not much different than those associated with tissue loss lesions. Microscopic lesions in normal fragments is not uncommon and has been documented elsewhere. For instance, in a coral disease survey from Micronesia [54] or New Caledonia [68], ca. 26% and 28%, respectively, of normal fragments had microscopic lesions comprising changes similar to those seen here. Fungi were the dominant organisms associated with tissue loss in Saudi Arabia in most species examined. In contrast, when organisms were associated with tissue loss lesions, ciliates dominated for *Acropora* in the Pacific [69] whereas chimeric parasitic corals dominated for *Montipora* in Hawaii [70]. Cell associated microbial aggregates (CAMAs) were associated with Pocilloporidae, Poritidae, and Acroporidae thus extending a pattern similar to that in the Indo-Pacific where CAMA infect these same coral families [31]. Presence of CAMA in *Stylophora* has not been previously described and adds another genus of Pocilloporidae to the list of members of this family infected with bacterial aggregates. This study also extends the presence of CAMAs in corals to the Red Sea (biogeographic range expansion).

#### Coral taxa differ in disease susceptibility

There were differences in disease prevalence among coral genera with *Acropora, Favites/Dipsastraea* and *Millepora* having a higher disease prevalence than expected based on their abundance in the field. This is consistent with other regions of the world where disease susceptibility differs across families or genera [40,55,64,71,72]. Our study differed from other regions, in that, over half of the coral genera within transects had signs of disease (16 out of 30 coral genera). In contrast, Aeby et al. [64] found seven out of 25 coral genera with disease signs in the Persian Gulf and Williams et al. [72] found five affected coral genera in the Line Islands which has approximately 31 coral genera on its reefs [73].

#### Regional differences in disease and potential environmental co-factors

The survey sites were spread out along a latitudinal gradient spanning from 19° to 24°N. Among the survey sites, coral cover (a measure of host abundance) ranged from 3% to 83%, degree of heat stress, as measured by DHW, ranged from 2.5 to 5.3 and amount of bleaching ranged from 0% to 54%. All three of these co-factors would be expected to affect subsequent disease prevalence and as expected, average disease prevalence varied from 0% to 1.9%. However, no significant relationship was found between disease prevalence and coral cover or DHW. And even more surprising, there was a negative relationship, not a positive one as expected, between percent coral bleaching and disease prevalence. This is in sharp contrast to what has been found on coral reefs in other regions. A positive relationship between host density and disease prevalence is considered a hallmark of the infectious process whereby higher host abundance results in greater rates of transmission and localized increases in prevalence [74]. In the Indo-Pacific, an association between coral cover and coral disease prevalence has been found in numerous regions [13,75–77]. Warmer temperatures, heat stress and bleaching have also been linked with higher disease prevalence [2,13,14,40]. For example, in the Persian Gulf, white syndrome outbreaks coincide with annual thermal heating events [78]. In the Caribbean, bleaching extent was linked to increased disease incidence [79] and tissue-loss disease outbreaks frequently follow bleaching events [80–82]. Our study suggests that a very different pattern is emerging for the Red Sea. The northern-most sites along the coast of Yanbu had the highest disease levels despite no bleaching occurring within transects and although heat stress was higher in this region, DHW alone was not a significant factor explaining disease prevalence.

Reef corals in the northern Red Sea have extraordinarily high thermal tolerances in relation to the ambient temperatures they usually experience [24] and our study supports this. Significant bleaching is expected when the DHW value reaches 4°C-weeks (https://www.coralreefwatch.noaa.gov/product/5km/index_5km_dhw.php) yet our sites in Yanbu had DHW values over 4 but no bleaching was observed. Thermal tolerance in corals has been linked to host factors [83–87], Symbiodiniaceae partners [88–90] or resident microbial communities [91]. In some coral species, thermal tolerance comes at the expense of increased disease susceptibility [92,93] and this has also been suggested as a possible explanation for high disease levels found in corals in the Persian Gulf [64]. Whether there are trade-offs between disease susceptibility and thermal tolerance in corals in the central Red Sea is a hypothesis worth exploring. No work as yet been done on latitudinal gradients of microbial communities or host adaptations on corals in the Red Sea. However, within the Red Sea, the main Symbiodiniaceae genus in *Porites* changed from *Durusdinium* (D1) at warmer nearshore location to *Cladocopium* (C15) at cooler offshore locations [94] suggesting that differences in Symbiodiniaceae could be influencing spatial patterns of disease occurrence in this region.

#### Disease prevalence is low despite environmental challenges

Compared to other ocean basins, the Red Sea experiences extreme temperature variation and regularly exceeds 32°C in the summers [18,19,95,96] which are conditions not well tolerated by most other corals. Chronic temperature stress can exert significant energetic costs on corals resulting in reduced growth and reproduction [97–99] and an increased prevalence of coral diseases [2,13]. Yet, we found corals existing in this challenging environment to have surprisingly low disease levels (<1%). Moreover, our surveys were conducted in the midst of a bleaching event. This suggests acclimation/adaptation to prevailing environmental conditions [19,100] and notably, the Red Sea is an arid region with minimal terrestrial run-off and almost no riverine input [18]. At our sites, we saw little evidence of sedimentation, water clarity was good and there was little macroalgae on the reef (<1%). Coastal coral reefs in other regions are increasingly exposed to excess nutrients, sediments, and pollutants discharged from land which are known to degrade local reefs [25] including increasing coral disease prevalence. Haapkyla et al. [11] found a 10-fold greater mean abundance of disease on reefs during the rainy summer months and concluded that rainfall and associated runoff were facilitating disease outbreaks. Laboratory studies showed that the rate of tissue loss from BBD increased with nutrient enrichment [26] and increased BBD prevalence in the field is associated with sewage effluent [101]. An experimental *in situ* nutrient enrichment of reefs was conducted in the Caribbean and corals exposed to chronic nutrient stress suffered a 3.5-fold increase in bleaching frequency and a two-fold increase in prevalence and severity of disease, compared to corals in control plots [27]. Terrestrial run-off also promotes the growth of macroalgae on coral reefs [25] which are major competitors with corals [102,103]. Additionally, macroalgae exude dissolved organic carbon which can disrupt the function of the coral holobiont and promote potential bacterial pathogens [104,105]. Although the coral reefs in the Red Sea have to contend with high temperatures and salinity, these appears to be countered by the lack of terrestrial run-off and all its associated problems. As a comparison, the Persian Gulf is also an arid region but has riverine input, and massive coastal habitat modification by dredging and converting shallow, productive marine areas into land for homes, recreation, and industrial activities [106]. Resuspension of sediments is an ongoing stress for coral reefs in this region as well [107]. Under these conditions, higher coral disease levels were found and attributed to environmental stress [108]. Compared to our study, reefs along the northeastern Arabian Peninsula show a 6-fold higher disease prevalence with a high number of localized disease outbreaks [64]. Reefs surrounding Kish Island, off the coast of Iran, showed a 20-fold increase in disease [109]. Disease is a serious problem in other world regions [3,12,40] and our study suggests that a reduction in human impacts and improvement in water quality may be effective management strategies giving corals increased capacity to withstand the warming oceans predicted with global climate change.

## Acknowledgments

Any use of trade, firm, or product names is for descriptive purposes only and does not imply endorsement by the US Government. We would like to thank the invaluable assistance of CMOR staff who helped with diving operations and boating.

